# GSAP Regulates Amyloid Beta Production through Modulation of Amyloid Precursor Protein Trafficking

**DOI:** 10.1101/2020.11.12.379313

**Authors:** Jerry C. Chang, Peng Xu, Eitan Wong, Marc Flajolet, Yue-Ming Li, Paul Greengard

## Abstract

In addition to participating in γ-secretase activity, presenilin 1 (PS1) regulates trafficking and subcellular localization of β-amyloid precursor protein (APP). We previously showed that gamma-secretase activating protein (GSAP) selectively modulates γ-secretase activity by inducing conformational change in PS1. However, little is known whether and how GSAP might influence APP trafficking and consequent generation of β-amyloid (Aβ) peptides. Here, to explore whether GSAP has any role in regulating APP trafficking, and to systematically investigate the intracellular trafficking routes of APP, we paired total internal reflection fluorescence microscopy, high-speed line scanning microscopy, and 4D microscopy with comprehensive imaging analysis methodologies to depict the elusive modes of APP trafficking at a single-vesicle level. Mobility and diffusivity changes reveal the existence of two kinetically distinct pathways, classified into mobile and immobile pools, for vesicular APP trafficking, suggesting high association between immobile vesicle pool and amyloidogenic processing. GSAP knockdown significantly lowers immobile pool without overturning APP vesicle diffusivity, suggesting that GSAP affects vesicular APP trafficking by retaining APP in membrane microdomains known to favor amyloidogenic processing. Our study reveals a novel role of GSAP in the regulation of Aβ-peptide formation that modulates switching of APP vesicles between immobile and mobile pools, which may help identifying new therapeutic strategies to treat Alzheimer’s disease.

## Introduction

One of the key neuropathological hallmarks of Alzheimer’s disease (AD) is the presence of extracellular senile plaques in the gray matter of the brain, which contain abundant β amyloid (Aβ)-peptide deposition^1^. Limiting Aβ-peptide production, without causing cell toxicity, is one of the primary AD therapeutic strategies^1–3^. Since Aβ peptides are generated from sequential proteolytic cleavage of β-amyloid precursor protein (APP), first by beta-site APP cleaving enzyme (BACE, β-secretase) and second by γ-secretase, vast resources have been used identifying and optimizing compounds that directly inhibit β- and γ-secretases. Unfortunately, all these inhibitors have failed clinical trials due to potential toxicity issues associated with altered cleavage of other important substrates, such as Notch. Recently, APP modifications and trafficking have been suggested as molecular mechanisms that drive AD pathogenesis ^4–6^. In agreement with this hypothesis, a large body of biochemical work has shown that intracellular trafficking and endosomal sorting of APP and β-cleaved carboxyl-terminal fragment of APP (APP-CTF, βCTF, C99) constitute APP-dependent pathogenic pathways in AD^7^. In addition, studies on late-onset Alzheimer’s disease (LOAD) also show that several LOAD risk factor genes encode proteins with known functions in regulating APP trafficking, including: PICALM^8^,^9^, SORL1^10–14^; SORCS1^14–16^, SORCS2^16^, SORCS3^16^, BIN1^17^ and CD2AP^17,18^. Furthermore, deletion/mutation of vesicular adaptor proteins that bind to APP, including Mint/X11 proteins^19^ and δ-COP^20^, alters APP cleavage and decreases Aβ-peptide generation in AD mouse models. These studies highlight modulation of APP trafficking as a potential therapeutic strategy for AD^21^.

Proteolytic processing of APP can be divided into non-amyloidogenic and amyloidogenic pathways. In general, 90% of APP enters the non-amyloidogenic pathway while only 10% undergoes amyloidogenic processing, and the regulation between the two is tightly correlated with APP trafficking^5^. The amyloidogenic processing of APP is known to be multifaceted, comprising of protein transport, cell surface localization and intracellular protein trafficking, and sequential cleavage by β- and γ-secretases^4–6^. Numerous studies indicate that the cleavage of APP to generate Aβ peptides occurs after APP is internalized into the endocytic pathway^22–24^. Despite the implications of amyloidogenic processing, research on the linkage between endosomal APP trafficking and Aβ-peptide generation remains challenging in living cells. However, recent developments in high-speed high-resolution microscopy platforms have enabled sufficient spatiotemporal resolution to begin to demonstrate how endosomal APP trafficking regulates Aβ generation in living cells. In neurons, it has been shown that APP is trafficked and cleaved in pre-synaptic vesicles^25–27^Using TIRF microscopy, Bauereiss *et al*. demonstrated that APP and BACE are exocytosed to the cell surface by distinct endosomal vesicles prior to the non-amyloidogenic cleavage^28^. In support of the functional role of APP in pre-synaptic vesicles, Yu *et al*. applied super-resolution microscopy and found that Aβ42-containing vesicles exclusively reside in pre-synapses and not in post-synapses^29^. Thus, identifying regulators associated with endosomal APP trafficking and Aβ-peptide generation with advanced microscopy and image analysis techniques are of primary importance.

We previously discovered a novel γ-secretase activating protein (GSAP) that participates in amyloidogenic processing without affecting Notch cleavage^30,31^. GSAP specifically regulates the γ-cleavage of APP through its interaction with the APP-CTF and presenilin 1 (PS1), the catalytic component of the γ-secretase complex^30^. Using an active-site–directed photoprobe, we demonstrated that the interaction between GSAP and PS1 favors γ-secretase to stay in its active conformation, which specifically increases the efficiency for γ-cleavage of APP without affecting Notch processing^31^. Mounting evidence suggests that PS1 also modulates Aβ production via regulating the sorting and trafficking of APP, and *vice versa^5^*. However, despite the influential effect of GSAP on modulating γ-secretase activity, whether GSAP plays any role in APP trafficking remains to be determined.

In the present study, we take a comprehensive imaging-based approach to address whether GSAP plays any role in modulating APP trafficking. Using total internal reflection fluorescence (TIRF) microscopy^32^ and custom-developed imaging analysis approach, we observed that a subset of surface APP vesicles, which are tethered near plasma membrane for seconds to tens of seconds (referred to as “immobile” pool), readily undergo membrane fusion and internalization processes. GSAP knockdown preferentially and significantly reduced the immobile pool of the surface APP vesicles, which potentially leads to a reduced level of Aβ-peptide release. In addition, combining high-speed line scanning confocal microscopy^33^ and a two-state kinetic modeling algorithm, we show that GSAP knockdown does not change the total number of cytoplasmic APP vesicles per cells. Instead, GSAP knockdown decreases the vesicle trajectory dwell time and reduces the probability of cytoplasmic APP vesicles resided in the BOUND state, suggesting a reduced cellular binding and association. Furthermore, we applied 4D microscopy for quantitative analysis of APP vesicle dynamics in whole cells. We demonstrated that GSAP knockdown effectively decreases the cellular constraints at longer timescales (> 100 seconds), which explains the reduced number of vesicles reaching cell surface. Finally, using density gradient centrifugation and immunoblotting analysis, we showed that GSAP preferentially accumulates in lipid rafts, where γ-secretase is known to reside for Aβ generation. Importantly, GSAP knockdown significantly reduced the level of APP-CTF in lipid rafts, suggesting that GSAP regulates amyloidogenic processing of APP via lipid raft-dependent pathways. Our study reveals a novel role of GSAP in Aβ formation that involves the regulation of APP trafficking.

## RESULTS

### GSAP knockdown decreases cell surface APP and Aβ peptide generation without altering total APP level

First, we determined the effect of GSAP knockdown on subcellular APP localization in N2a cells. In order to better visualize the abundance of intracellular APP pool, we created a stable APP-GFP overexpressing N2a cell line (N2a-695GFP), which constitutively expresses C-terminal fusion of GFP to full-length APP. This is a well-tested labeling method known to accurately preserve the distribution and localization of full-length APP without altering APP trafficking^34,35^. A sequence-specific siRNA was applied to knockdown GSAP expression in N2a-695GFP cells, which effectively decreases Aβ production (Fig. 1a) and had no sign of apparent side effects on cell viability and proliferation (Suppl. Figure 1-3). We also confirmed by western-blot analysis that siRNA-induced GSAP knockdown does not alter the expression level of full-length APP in N2a-695 cells (Fig. 1b). To validate the distribution of APP in N2a-695GFP cells, we performed live-cell confocal microscopy. We found that the majority of APP-GFP is located in an organellar pool residing within the Golgi complex (Fig. 1c, right), which is consistent with previous findings^4,35^. Interestingly, GSAP knockdown does not significantly alter the major cellular APP-GFP allocation at steady state (Fig. 1c, left), as both control and GSAP knockdown cells show no significant difference in regarding to how APP colocalized with the Golgi apparatus (Fig. 1d).

**Figure 1.**
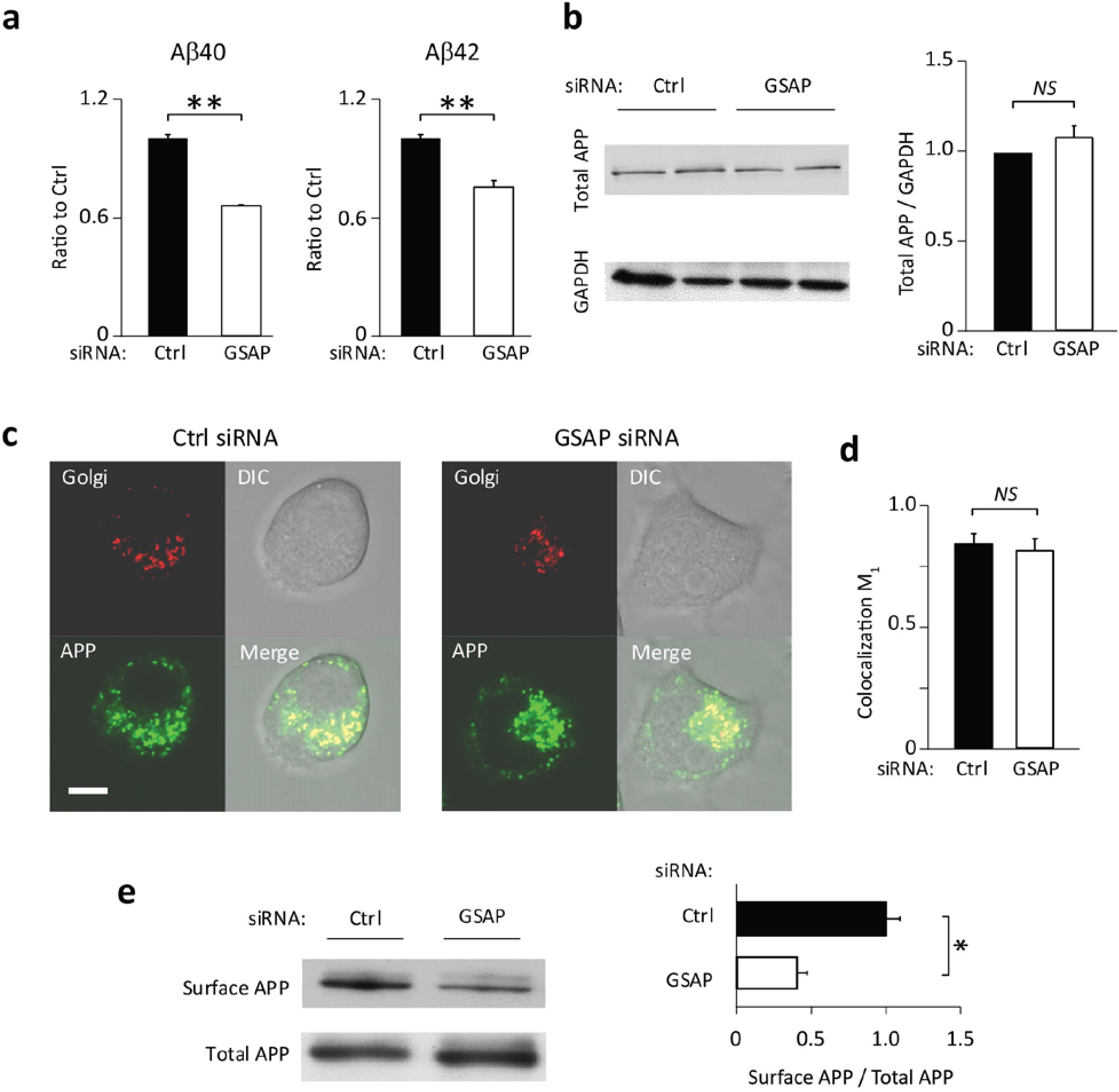
Effect of GSAP knockdown on Aβ secretation and APP distribution. (a) Downregulation of Aβ40 and Aβ42 secretation by GSAP knockdown using control (CTRL) and GSAP siRNA in N2a-695 cells. Secreted Aβ40 and Aβ42 levels were measured by ELISA (mean±SEM; n=3). (b) Immunoblotting analysis of the total APP protein level following transfection with CTRL and GSAP siRNA (Left) and quantification (Right). (mean±SEM; n=5). (c) Representative images of subcellular localization of APP and Golgi in living N2a cells infected with a baculovirus expressing an RFP-Golgi marker, and then transfected with APP-GFP plasmids. (d) Colocalization coefficients m1 quantification for APP-GFP and RFP-Golgi (n = 12; Scale bar: 10 μm). (e) Cell surface APP was biotinylated and precipitated with streptavidin-coupled beads following transfection with CTRL and GSAP siRNA. Total APP in the lysates and the precipitated surface APP were detected by immunoblotting (Left) and quantification (Right) (means±SEM; n=3). *P<0.05; **P<0.01; ^NS^p > 0.05.

In addition to the APP pool in trans-Golgi network (TGN), surface APP is also known to participate in Aβ production^36^ so we determined the effect of GSAP knockdown on the level of cell surface APP. Remarkably, using surface biotinylation we found that GSAP knockdown drastically reduces the levels of surface APP (Fig. 1e). Since surface APP only counts for 10 percent of total APP in the cells^4^, it was not possible to image surface pool of APP-GFP in living N2a cells using confocal microscopy due to fundamental sensitivity limitations (Fig. 1c). Nevertheless, these results strongly suggest that GSAP may play a role in modulating surface APP trafficking.

### Imaging single APP vesicles at the cell surface

Previous imaging studies have shown that expression of cell surface APP is concentrated into punctate structures^28,37,38^, which are associated with clathrin-coated vesicles^39^. To detect cell surface APP-GFP with high spatial and temporal resolution, we custom-built a total internal reflection fluorescence (TIRF) microscope, which offers single-photon sensitivity at full frame rates. In accordance with previously reported studies, our TIRF imaging showed that GFP-tagged cell surface APP demonstrates punctate structures resembling vesicles observed in the living N2a695-GFP cells (Fig. 2a, see also Suppl. Video 1). We noticed that the around 80% of single vesicle trajectories exhibit short displacements consistent with “Confined” diffusion (Fig. 2b, right), and the remaining trajectories appear visually “Diffusive” (Fig. 2b, left). Based on their vertical movements towards or away from the plasma membrane, the motion patterns of the confined APP-GFP vesicles were further classified into three groups (Fig. 2c): “Tethering” events showed confined APP-GFP vesicles continuously appearing in the TIRF field (within ≤100 nm^40^) and the central positions of the fluorescent signal remained stationary for a few seconds up to tens of seconds; “Fusion” events showed that fluorescent signal of these stationary vesicles vanishes abruptly and completely, indicating spreading of the APP-GFP into the plasma membrane upon membrane fusion; “Internalization” events showed the sudden appearance of the fluorescent signal that stays stationary for a few seconds up to tens of seconds in the TIRF field, indicating the recruitment of newly tethered vesicles from the plasma membrane. To quantitatively characterize the overall motions of single APP-GFP vesicles at the local environments close to plasma membrane, we calculated the mean-squared displacement (MSD, 〈*γ^2^*〉) in function of incrementing time intervals, a classical model for studying the diffusivity of single particles^41,42^. In brief, MSD increases linearly with time when particles undergo Brownian motion; whereas MSD versus time rapidly reaches a plateau level when particles evolve in crowded or confined environments^41,42^. We found that MSD versus time scatter plot showed the best fit (light blue curve, *R^2^* = 0.9782) to an exponential rise to a maximum function (Fig. 2d), demonstrating that the movements of single APP-GFP vesicles near plasma membrane becomes apparently restricted, which are mostly due to constraints from neighboring obstruction^43,44^.

**Figure 2.**
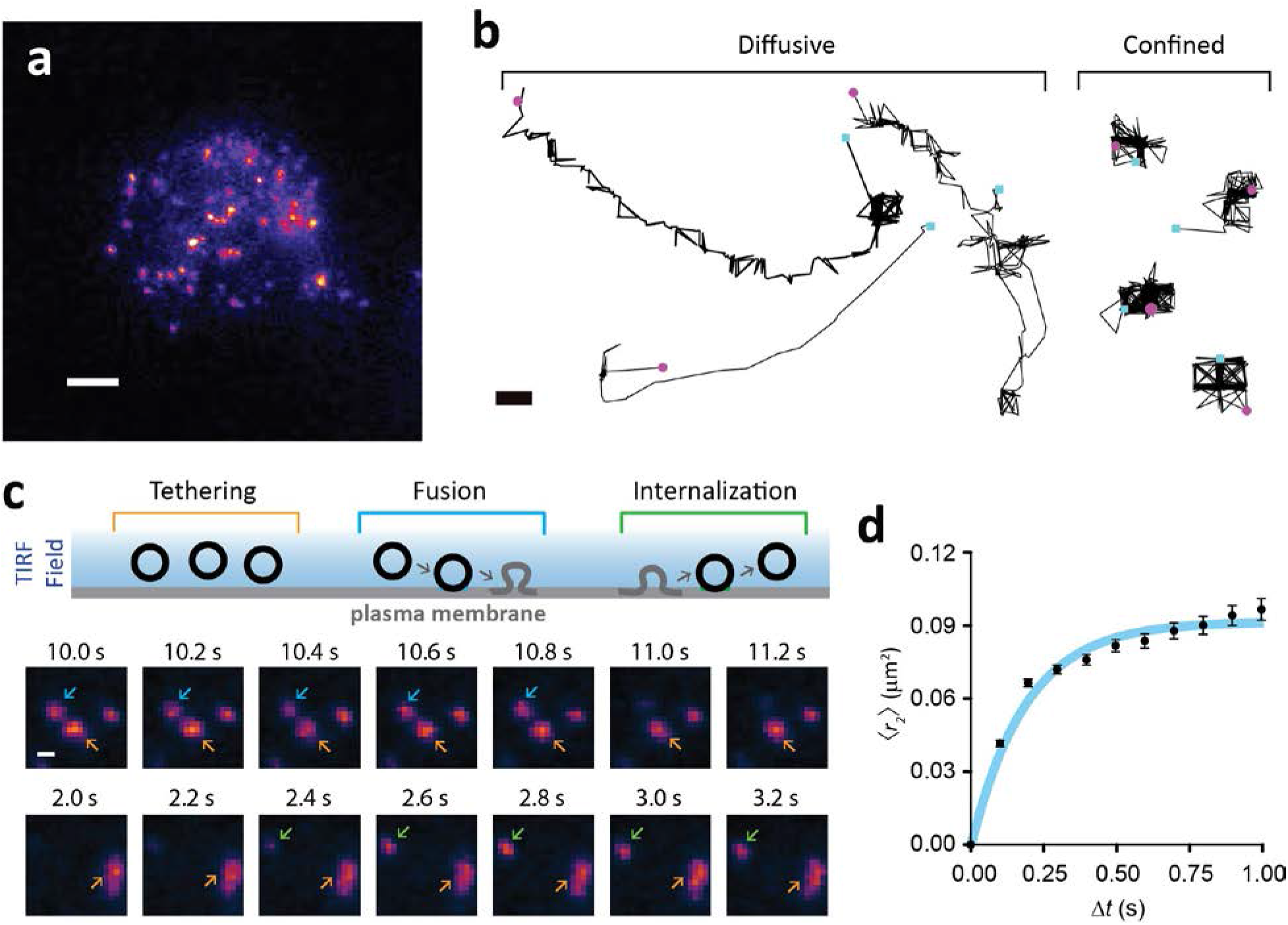
Tracking dynamic movement of surface APP vesicles in living cells. (a) The first frame from a time-lapse TIRF recording shows the surface distribution of APP-GFP in live N2a695GFP cells (see suppl. Video 1 for complete 60 s time-lapse recording). Signals of APP-GFP vesicles are displayed with ImageJ lookup table ‘Fire’. Scale bar: 5 μm. (b) Schematic diagrams and representative sequential time-lapse images of the fluorescent changes reflect 3 major diffusive patterns of surface APP-GFP vesicles. (1) Tethering: vesicles near surface remain stationary over time; (2) Fusion: tethered vesicles fuse with the plasma membrane and lose fluorescent signal; (3) Internalization: vesicles formed from the plasma membrane then get internalized and join the tethered vesicle pool. Examples of sequential images of vesicle “Tethering”, “Fusion”, and “Internalization” events are indicated by orange, blue, and green arrows, respectively. Images were recorded over time at a 10 frames s^−1^ rate and displayed here a reduced 5 frames s^−1^ rate for simplicity. (c) Representative trajectories of two types of movement were observed: *Confined* and *Diffusive* (see text). The start points (cyan square) and ends point (purple cycle) within each trajectory are shown. Scale bar: 200 nm. (d) The averaged MSD vs time plot of surface APP-GFP vesicles. Blue curve in the plot represents an exponential rise to a maximum fit (*R^2^* = 0.9782). The result is presented as averages ± SEM of 11 cells from 3 independent experiments.

To systematically profile the motion confinements of APP-GFP vesicles close to plasma membrane, we developed a trajectory analysis algorithm that effectively classifies the immobile and mobile particles in a robust and computationally efficient fashion. The rationale behind our approach is that the average intensity projection (AIP) of time-lapse images, which averages all the pixel values in the time-series stacks, directly permits the visualization of immobile particles (Fig. 3a). As such, the initial trajectory location of one immobile particle and the centroid of the fluorescent spot at the AIP image generated by the exact same immobile particle should be within proximity. The proposed method is achieved in three steps: (1) spot detection, (2) local vicinity analysis, and (3) immobile versus mobile vesicle classification (Fig. 3b). Representative trajectories of single APP vesicles in a N2a-695GFP cell obtained by single particle tracking of time-lapse TIRF imaging, segregated based on the immobile/mobile classification, are shown in Figure 3c. Based on these results, approximately one fifth of the trajectories were characterized as immobile (19.37 %, *n* = 581) while the remaining trajectories were classified as mobile (80.63 %, *n* = 2484). In addition, single particle diffusion rates (*D*) of the immobile and mobile pools are significantly different (*\*\*\*p* < 0.001, Mann-Whitney *U* Test) (Fig. 3d). However, both pools of trajectories displayed heterogeneous diffusivity (Immobile: *D_medium_* = 0.0193 μm^2^/s, IQR 0.0170 - 0.0233; Mobile: *D_medium_* = 0.0353 μm^2^/s, IQR 0.0278 - 0.0460). We note a large overlap of diffusion rates between immobile and mobile pools (Fig. 3d), indicating the occurrence of the intermediate states in between.

**Figure 3.**
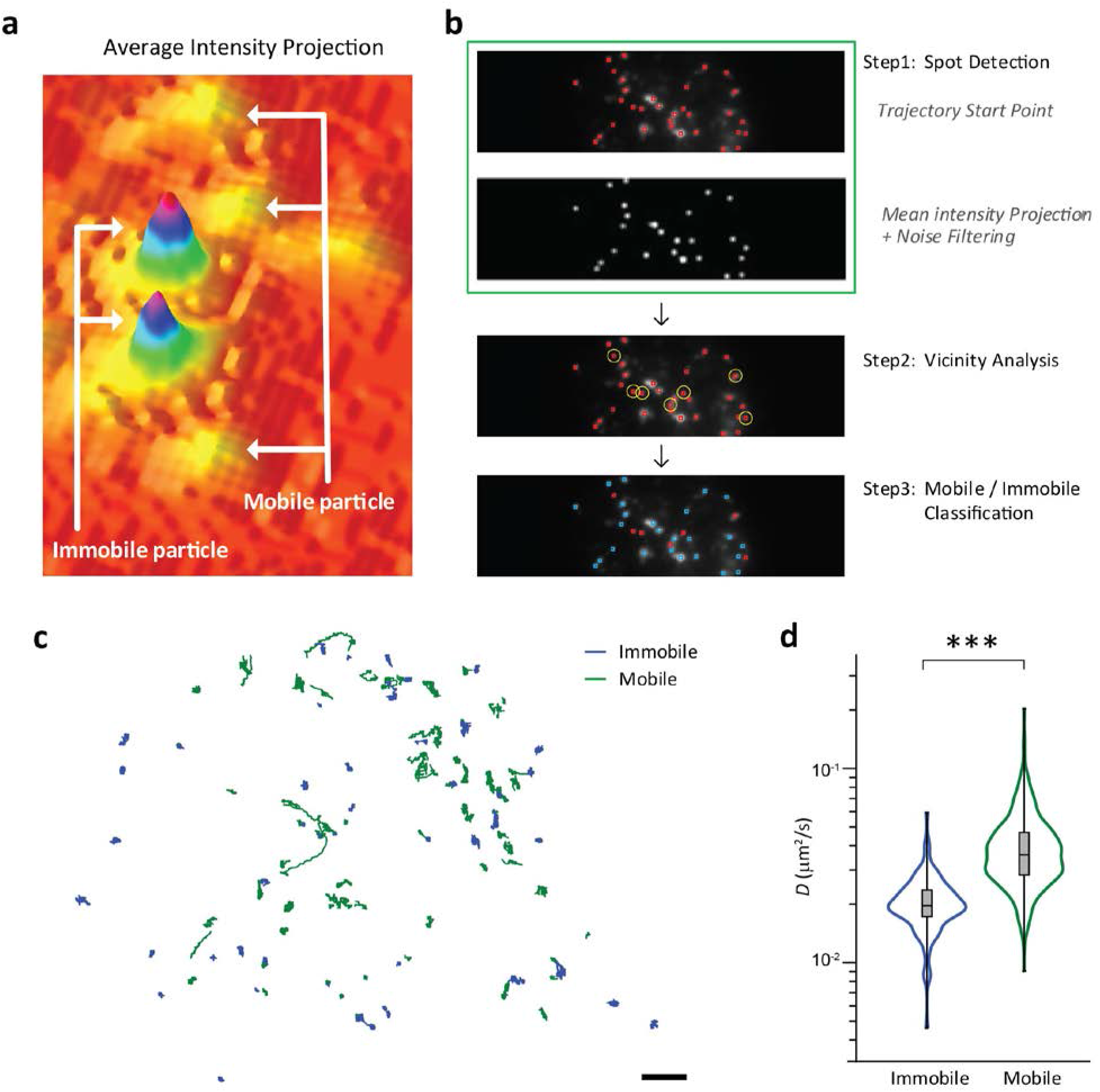
Classification of immobile and mobile surface APP-GFP vesicles. (a) Schematic illustrating the immobile and mobile vesicles projected on an Average Intensity Projection (AIP) of a time-lapse TIRF image sequence. The intensity from the mobile particles is averaged out with background noise due to continuous movement over time or transient appearance, while immobile particles show limited movement over longer time are easily identified on AIP. (b) Overview of the image processing pipeline of immobile/mobile vesicle classification. The processing steps include (1) Spot Detection: The red squares displayed in the top panel shows the starting points of all the trajectories identified in a time-lapse TIRF recording. The bottom panel shows the AIP of the same time-lapse TIRF recording. To make the center of immobile particles easily segmented, AIP image is further subjected for noise filtering; (2) Vicinity Analysis: using the nearest-neighbor method, if the distance from the starting point of one trajectory to the closest spot depicted from the AIP is above the mean trajectory displacement value (~ 250 nm), the starting point is circulated in yellow. The script runs in loop until the end; (3) Mobile/Immobile classification: all the trajectories identified by the Vicinity Analysis from step 2 (starting points marked in red squares) are in the Mobile pool, all the other trajectories (starting points marked in blue squares) are in the Immobile pool. (c) The representative trajectory maps of surface APP-GFP vesicles acquired by TIRF imaging revealed Mobile and Immobile pools of surface trafficking. (d) Comparison on the distribution of diffusion coefficients of Mobile and Immobile pools of single APP-GFP vesicles. Boxplots represent mediums and one-sigma confidence intervals (rectangles). The distributions were compared using the Mann-Whitney *U* test. ***p < 0.001.

### GSAP knockdown reduces the number of APP vesicles with elevated diffusivity

The limited restriction of vesicle mobility close to plasma membrane led us to wonder whether GSAP knockdown specifically alters the trafficking of immobile and mobile pools of APP-GFP vesicles, or both. We applied the immobile/mobile classification algorithm to compare the difference between the trajectories subjected to either control or GSAP siRNA treatments. The quantification of the immobile pool of APP trajectories per cell revealed a drastic decrease (18.97 %, *\*\*p* = 0.00361, two-tailed test) with GSAP knockdown (Fig. 4a Left). However, when considering the diffusion rates in the immobile pool, we did not find a significant difference (*p* = 0.1, Mann-Whitney *U* Test) between cells treated with control and GSAP siRNA (Fig. 4a Right), suggesting that GSAP knockdown does not alter the stability of the microenvironments where the immobile vesicles reside. In addition, treatment with GSAP siRNA caused a modest but statistically insignificant (*p* = 0.157, Mann-Whitney *U* Test) decrease of APP trajectories per cell in the mobile pool as compared to control (Fig. 4b Left). Furthermore, GSAP knockdown led to a significant (\**p* = 0.013, Mann-Whitney *U* Test), but slight, increase in diffusion rates of the mobile pool as compared to control (Control: *D_medium_* = 0.0355 μm^2^/s, IQR 0.0262 - 0.0463, *n* = 4229; GSAP knockdown: *D_medium_* = 0.0372 μm^2^/s, IQR 0.0281 - 0.0491, *n* = 4203, τ cutoff > 2 s) (Fig. 4b Right), possibly reflecting the mild reduction of trajectories in the mobile pool after GSAP knockdown (Fig. 4b Left). Regardless, the total trajectories per cell is significantly reduced after GSAP knockdown (\**p* = 0.046, Mann-Whitney *U* Test) (Fig. 4c). This is consistent with the surface biotinylation data in Figure 1d, for which a significant decrease in surface APP level was found after GSAP knockdown.

**Figure 4.**
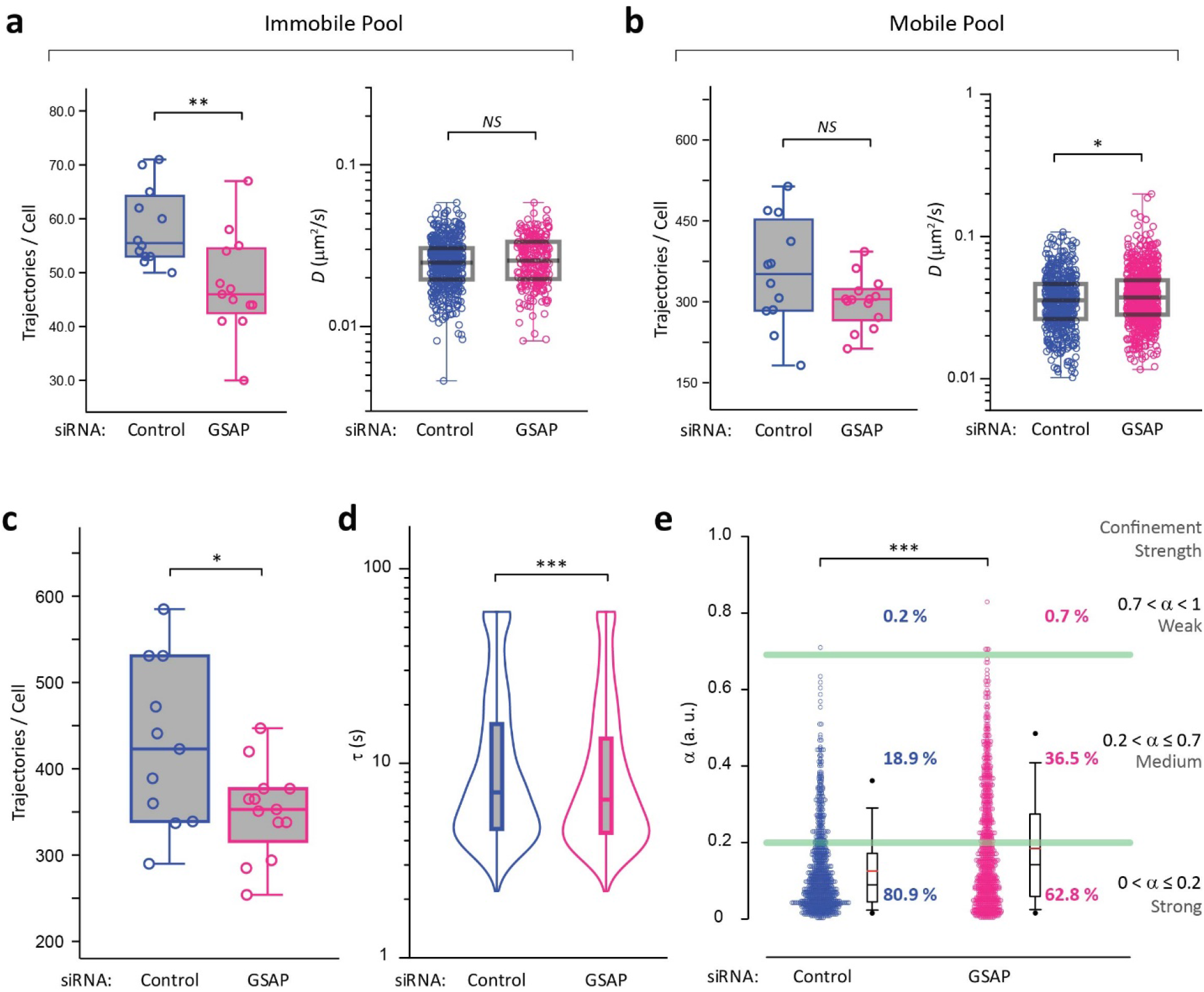
GSAP modulates confinement of surface APP vesicles. (a) Trajectories per cell and diffusion coefficient distribution from the immobile pool of trajectories subjected to control or GSAP siRNA treatments. (a) Trajectories per cell and diffusion coefficient distribution from mobile pool of trajectories subjected to control or GSAP siRNA treatments. (c) Trajectories per cell subjected to control or GSAP siRNA treatments. (d) Comparison on the trajectory dwell time of single APP-GFP vesicles subjected to control or GSAP siRNA treatments. (e) Comparison on the diffusivity of single APP-GFP vesicles subjected to control or GSAP siRNA treatments. Classification of diffusion model is based on the diffusional anomality α value: normal diffusion (1 ≥ α > 0.7), partially corralled diffusion (0.7 ≥ α > 0.2), and confined diffusion (0.2 ≥α > 0). Note that no super diffusion (α > 1) was found in our analysis. A significant increase of the α value distribtion in the GSAP knockdown group, indicating the effect of GSAP on the diffusional confinement of surface APP vesicles. Comparison on the distribution of diffusion coefficients of Mobile and Immobile pools of single APP-GFP vesicles. Boxplots represent mediums and one-sigma confidence intervals (rectangles). The distributions were compared using the Mann-Whitney *U* test. ***p < 0.001; **p < 0.01; *p < 0.05; ^NS^p > 0.05.

Reduction of the APP vesicles in the immobile pool after GSAP knockdown was further confirmed by an analysis of the trajectory dwell time (τ). In principal, the higher ratio of APP vesicles stays in the mobile pool, the higher probability of short trajectory dwell times is detected. As expected, significantly higher levels of short dwell times (Control: τ_medium_ = 7.1 IQR 4.6 - 15.9, *n* = 2322; GSAP knockdown: τ_medium_ = 6.5 s, IQR 4.4 – 13.4, n= 2443; \*\*\**p* <0.001, Mann-Whitney *U* Test) were found in the trajectories treated with GSAP siRNA as compared to control (Fig. 4d). In order to obtain more insight in the diffusion pattern of surface APP vesicles, we calculated the diffusivity of individual trajectories based on the anomalous diffusion equation ^41^:

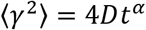
where 〈*γ^2^*〉 is the mean-squared displacement, *D* is the diffusion coefficient, *t* is the elapsed time, and *α* is the sub-diffusive exponent. In case of α > 1, α = 1, and α < 1, the particle undergoes super-diffusion, normal diffusion (Brownian motion), and sub-diffusion, respectively. According to this analysis, all the surface APP vesicles exhibit sub-diffusive lateral motion (α < 1), and neither super-diffusion (α > 1) nor normal diffusion (α = 1) were identified in both control and GSAP siRNA treated cells. In cells treated with control siRNA, APP vesicles diffuse slowly at *D* = 0.0402 ± 0.0183 μm^2^/s (mean ± SD, *n* = 1707, τ cutoff > 2 s). In contrast, GSAP knockdown led to a significant increase (\*\*\**p* < 0.001, Mann-Whitney *U* Test) in APP vesicle diffusivity (*D* = 0.0479 ± 0.0255 μm^2^/s, Mean ± SD, *n* = 1401, τ cutoff > 2 s). We took advantage of the *α* value measurement to further classify the variations in confinement strength into three states, namely weak confinement (1 > *α* > 0.7), medium confinement (0.7 ≥ *α* > 0.2), and strong confinement (0.2 ≥ *α* > 0). In the control cells, we found 80.9 %, 18.9%, and 18.9% of vesicles corresponding to weak, medium, and strong confinement, respectively (Fig. 4e). In contrast, in the GSAP knockdown cells, 62.8%, 36.5% and 0.7% of APP vesicles were corresponding to weak, medium, and strong confinement, respectively (Fig. 4e). Interestingly, the percentage of medium confinement subgroup is nearly doubled after GSAP depletion (18.9 % *vs*. 36.5 %). To summarize, our observations suggest that the effect of GSAP knockdown on surface APP trafficking is likely due to releasing a subset of immobile APP vesicles away from microenvironmental constraints near plasma membrane, which results in an increase of sub-diffusive exponent *α*.

### GSAP regulates intracellular APP trafficking

Since GSAP knockdown lowers the probability of APP vesicles residing in the immobile pool, next we asked whether the effect of GSAP knockdown on the motion of APP vesicles is solely reserved for APP vesicles near the plasma membrane. To address this specific question, we employed an ultrafast line-scanning confocal system, which provides a high-speed image acquisition in excess of one hundred frames per second at full resolution,^45^ for imaging the motion of intracellular APP vesicles in real time. As shown in Figure 1c, intracellular APP-GFP predominantly co-localized with the Golgi marker. As such, our imaging focal plane was set at a vertical level between Golgi and the plasma membrane (about 2-3 μm above the coverslip), where the single APP-GFP vesicles were easily captured with minimal interference from the Golgi apparatus (Fig. 5a, see also suppl. Figure 4). To monitor the lateral movement of intracellular APP-GFP vesicles with sub-pixel accuracy, we limited our 2D imaging at 10 Hz to provide sufficiently high signal-to-noise ratio (SNR ≥ 5) for SPT ^46–48^. Visual comparison of the trajectories showed that N2a-695GSP cells subjected to control siRNA are overall seen in confined space (Fig. 5b top; see also Suppl. Video 2), whereas cells treated with GSAP siRNA appear to be more diffusive (Fig. 5b bottom; see also Suppl. Video 3). We then calculated the displacement (Δd) of each trajectory, and we found a significant increase in displacement between the groups of control and GSAP knockdown (Fig. 5c; Control: Δd_medium_ = 0.284 μm, IQR 0.168 – 0.545, *n* = 1890; GSAP knockdown: Δd_medium_ = 0.414 μm, IQR 0.229 – 0.765, *n* = 1913; τ cutoff > 2 s; \*\*\**p* < 0.001, Mann-Whitney *U* Test). In addition, we evaluated the dwell time for each trajectory and found a significant decrease in trajectory dwell times after GSAP knockdown as compared to control (Fig. 5d; Control: τ_medium_ = 4.4 s, IQR 2.375 – 13.225, n = 1890; GSAP knockdown: τ_medium_ = 3.8 s, IQR 2.2 – 9.5, n = 1913; τ cutoff > 2 s; \*\*\**p* < 0.001, Mann-Whitney *U* Test). This result is consistent with our trajectory dwell time data using TIRF imaging (Fig. 4d) where the trajectories subjected to GSAP siRNA treatment tend to display shorter dwell times. However, unlike data obtained from TIRF imaging (Fig. 4c), we do not find a significant difference (*p* < 0.650, Mann-Whitney *U* Test) in mean trajectories per cell between the control and GSAP knockdown cells (Fig. 5e). Given no apparent differences in the number of intracellular APP vesicles, our data suggest that the effects of GSAP knockdown on intracellular APP vesicles should have minimal impact on APP vesicle formation and release, and more likely impact vesicle motion, which is regulated by intracellular machinery responsible for the vesicle clustering and transport^49,50^.

**Figure 5.**
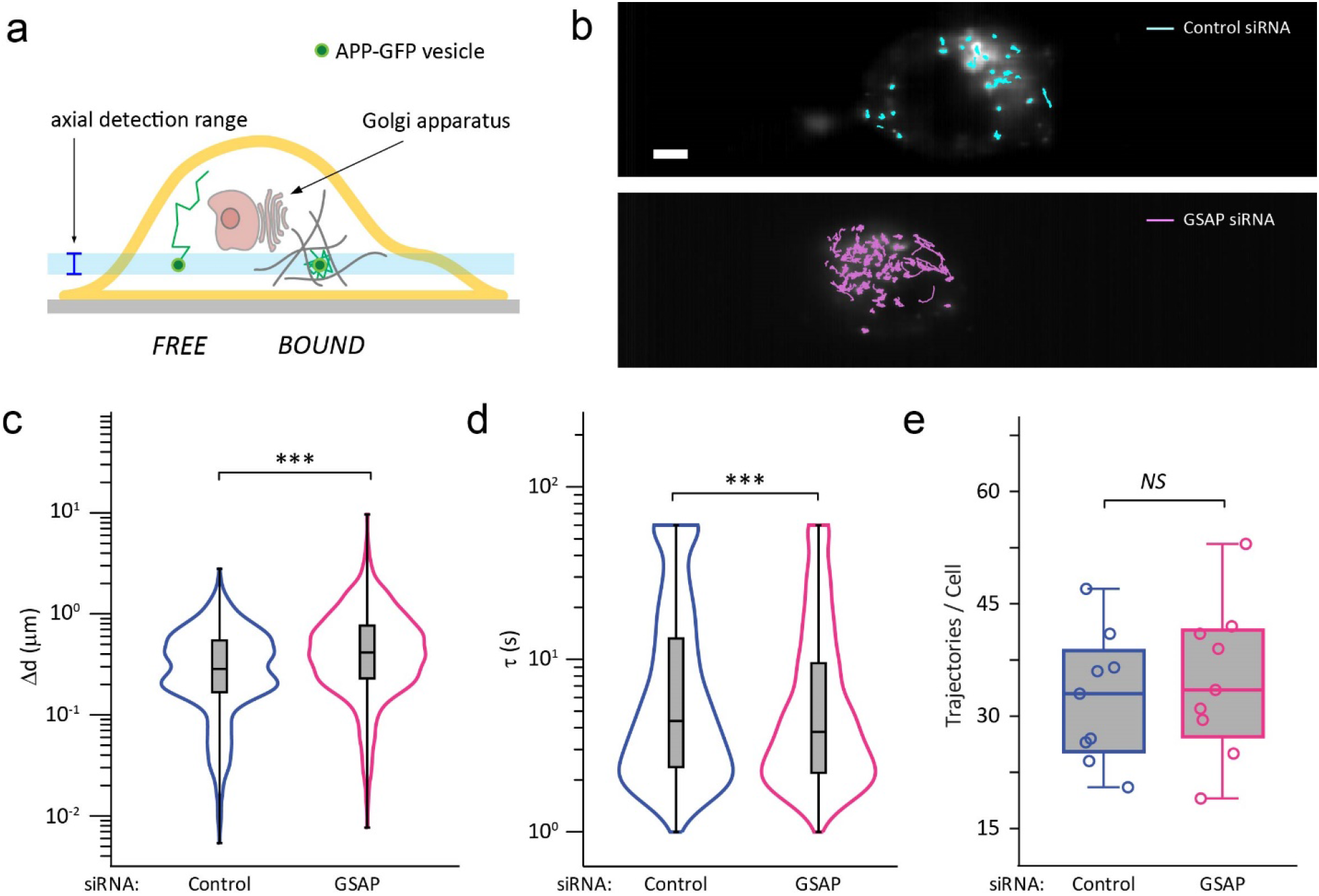
GSAP modulates vascular APP trafficking within the cytoplasmic environment. (a) Schematic of live-cell line-scanning microscopy showing the axial detection range that allows detection of single APP-GFP vesicles within the cytoplasmic environment in the proximity between Golgi apparatus and the plasma membrane. Vesicles without and with neighboring constraints are noted as *FREE* and *BOUND*, respectively. (b) Representative average intensity projection images superimposed with trajectories of single APP vesicles in N2a-695GFP cells subjected to control or GSAP siRNA treatments. Images were recorded at 10Hz for 60 seconds. The trajectory lifetimes shorter than one second were filtered and excluded. Scale bar: 5 μm. (c) Comparison of the distribution of trajectory displacement (Δd) single APP vesicles subjected to control siRNA and GSAP siRNA treatments. (d) Violin plots show the trajectory lifetime distributions of single APP vesicles subjected to control siRNA and GSAP siRNA treatments. (c) Comparison on the mean trajectories per cell subjected to control siRNA and GSAP siRNA treatments. Control and GSAP siRNA treated groups were collected from 3 independent experiments with 1890 and 1912 trajectories, respectively. Boxplots show mediums and one-sigma confidence intervals (rectangles). The distributions were compared using the Mann-Whitney *U* test (c and d) or two-tailed test (e). ***p < 0.001; ^NS^p > 0.05.

To better understand the partitioning and trafficking of APP vesicles, we fit a two-state kinetic modeling algorithm, termed Spot-On^51^, which separates heterogeneous diffusivity of single particle motions into either a BOUND state, characterized by slow diffusion, or a FREE state, associated with fast diffusion. In addition, Spot-On offers two ways to probe the diffusional motion. In the single-cell analysis, FREE state diffusivity (*D_FREE_*), BOUND state diffusivity (*D_BOUND_*), and BOUND fraction are extracted from each cell with hundreds to thousands of trajectories. In the global analysis, tens of thousands of trajectories across multiple cells are combined to directly fit the two-state kinetic model for acquiring the *D_FREE_, D_BOUND_* and BOUND fraction. We first applied the single-cell analysis to classify trajectories into BOUND and FREE states (Control: 12 cells; GSAP siRNA: 16 cells; 1000-3000 trajectories/cell, no τ cutoff). Figure 6a shows representative examples of model-fitting displacement histograms of trajectories from a single cell subjected to either control (Left) or GSAP (Right) siRNA treatments, where the GSAP siRNA treated cell displayed long-tail distributions over time, indicating an increase of FREE state. Further classification of BOUND and FREE states based on single-cell analysis observe no significant differences in *D_BOUND_* (Fig. 6b) and *D_FREE_* (Fig. 6c) between control and GSAP siRNA treated cells. However, the fraction of single APP vesicles in the BOUND state decreased significantly after GSAP siRNA treatments (Fig. 6d). In agreement with results obtained from single-cell analysis, global trajectory analysis from the ensemble trajectories found similar fraction of BOUND state with no difference in *D_BOUND_* (Table 1). However, unlike the single-cell analysis, the global trajectory analysis show that control and GSAP siRNA treated groups were statistically significant for *D_FREE_* (Table 1). Nevertheless, taken together, the results strongly suggest that the effect of GASP knockdown on intracellular APP trafficking increases the probability of state changes of APP vesicles from BOUND state to FREE state. Note that diffusion results obtained from intracellular vesicle tracking shows a good agreement with results from surface vesicle tracking (Fig. 4), indicating that the modulation of APP trafficking by GSAP contributes to both surface and intracellular APP vesicles.

**Figure 6.**
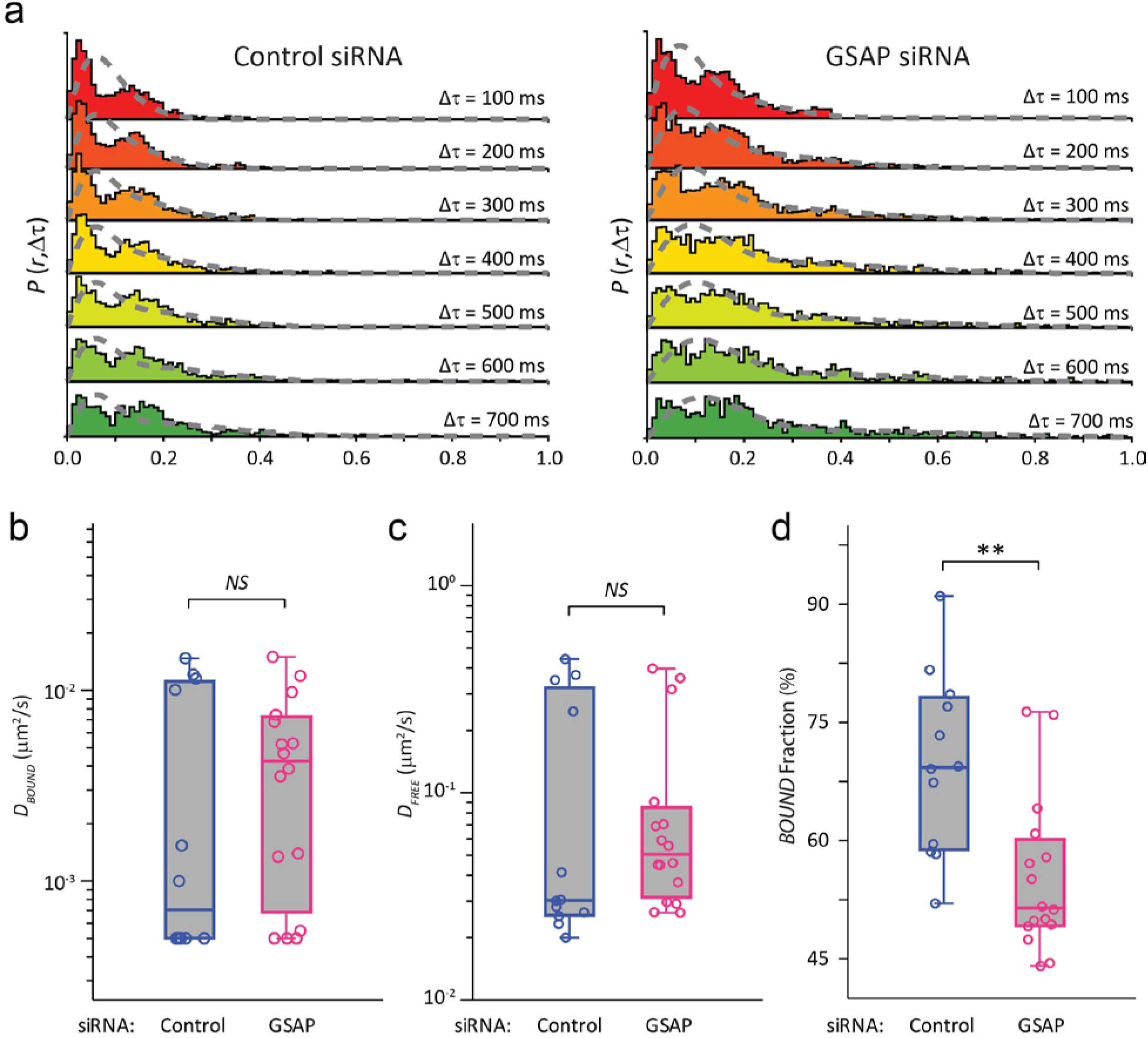
GSAP knockdown decreases cytoplasmic APP vesicles in the *BOUND* state. (a) Representative displacement histograms for trajectories of APP vesicles in N2a-695GFP cells. The jump lengths distribution is displayed in consecutive time frames, from Δτ = 100 ms to Δτ = 700 ms. Control siRNA treated group is shown of the left panel and GSAP siRNA treated group is shown on the right panel. The distributions in the GSAP siRNA treated cells show a longer tail indicative of a larger *FREE* population. Gray dash lines are fits from two-state kinetic modelling. (b) Quantification of diffusion coefficients of APP-GFP vesicles in *BOUND* fraction shows no statistically significant difference between cells subjected to control siRNA and GSAP siRNA treatments (p = 0.471). (c) Quantification of diffusion coefficients of APP-GFP vesicles in *FREE* fraction shows no statistically significant difference between cells subjected to control siRNA and GSAP siRNA treatments (p = 0.318). (d) Quantification of trajectories in the *BOUND* fraction shows a significant decrease in the cells subjected to GSAP siRNA treatments as compared to control (p = 0.00138). Boxplots show mediums and one-sigma confidence intervals (rectangles). The distributions were compared using two-tailed test. **p < 0.01; ^NS^p > 0.05.

**Table 1.** Statistics of *BOUND* and *FREE* states of the cytoplasmic APP-GFP vesicles. Fraction of *BOUND* vesicles and average diffusion coefficient of the *BOUND* and *FREE* vesicles are determined in either an average of individual cells (Single-Cell Analysis) or distribution of the total populations of single vesicle trajectories (Global Trajectory Analysis). Single-Cell Analysis: Control siRNA, *n* = 12 cells; GSAP siRNA, *n* = 14 cells. Global Trajectory Analysis: Control siRNA, *n* = 17356 trajectories; GSAP siRNA, *n* = 21617 trajectories. Data were collected from three independent experiments. The p-values are calculated from two-tailed test. ****p < 0.0001; ***p < 0.001; **p < 0.01; *p < 0.05.

| Single-Cell Analysis |  |  |  |
| --- | --- | --- | --- |
|  | Control siRNA | GSAP siRNA | Significance |
| $D_{BOUND}$ (mean ± SD) | $0.0045 \pm 0.00572$ | $0.00489 \pm 0.00439$ | $p = 0.471$ |
| $D_{FREE}$ (mean ± SD) | $0.136 \pm 0.165$ | $0.106 \pm 0.137$ | $p = 0.318$ |
| Bound Fraction (%) | $69.7 \pm 11.4$ | $55.3 \pm 9.84$ | $**p = 0.00138$ |
| Global Trajectory Analysis |  |  |  |
|  | Control siRNA | GSAP siRNA | Significance |
| $D_{BOUND}$ (mean ± SD) | $0.001 \pm 0.0001$ | $0.001 \pm 0.0002$ | $p = 1$ |
| $D_{FREE}$ (mean ± SD) | $0.031 \pm 0.0004$ | $0.050 \pm 0.0005$ | $****p < 0.0001$ |
| Bound Fraction (%) | $64.6 \pm 0.48$ | $55.5 \pm 0.33$ | $****p < 0.0001$ |

### Quantitative 4D Image Analysis of Spatiotemporal Control of APP trafficking through GSAP

We next sought to evaluate the effect of GSAP knockdown on the motion of APP vesicles in the entire cell volume using 3D time-lapse live-cell imaging, referred to as 4D microscopy^52^. To quantitatively analyze the SPT data generated by 4D microscopy with sufficient spatiotemporal resolution, the detected z range was limited to 19 μm with step-size of 0.5 μm, which resulted in a sequence-repeat 6-sec interval throughout the entire z steps. However, due to the repeated illumination of the entire imaging volume applied in 4D microscopy, which, unfortunately, caused unavoidable photobleaching. Importantly, compensating for that, acquisition time was limited to 10 minutes, and 80 to 200 trajectories were extracted from each set of time-lapse z-stack data using Imaris 8.4 (Bitplane). We also noted that by applying the upper and lower threshold of the “Quality”, which reflects the central intensity of the spot being detected, single APP-GFP vesicles were detected with the Imaris software without collecting high-density labeling of APP in the Golgi. Figure 7a (see also Suppl. Video 4 and 5) shows representative images of time-color-coded trajectories of single APP-GFP vesicles superimposed on the 3D projected image of a single N2a-695GFP cell, subjected to either control (top) or GSAP (bottom) siRNA treatments. In short, this approach allows for tracking the motion patterns of APP at a single vesicle level in 3D over extended periods of time.

**Figure 7.**
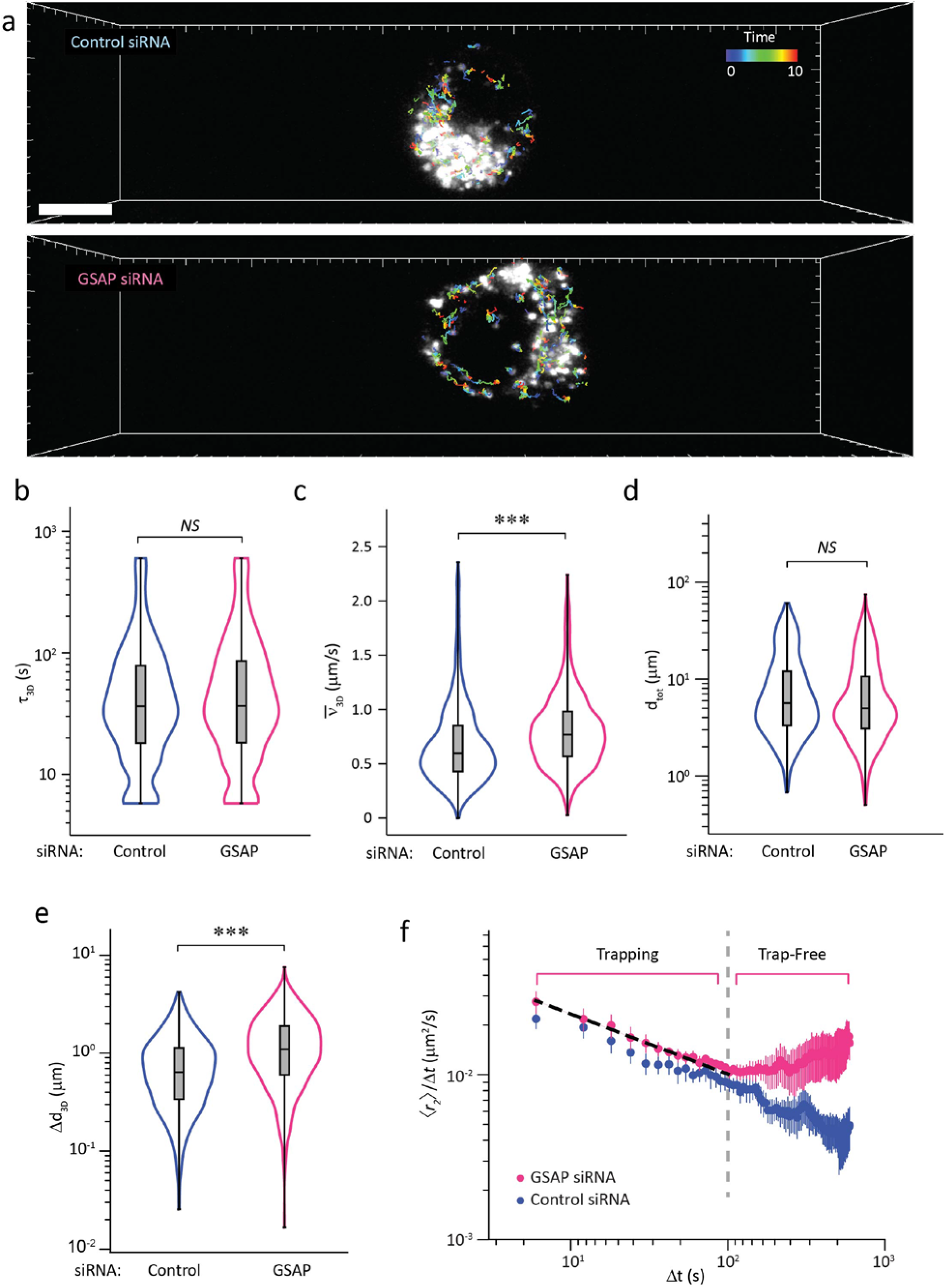
GSAP knockdown effects on anomalous subdiffusion of single APP vesicles revealed by 4D microscopy. (a) Representative three-dimensional (3D) images of single N2a-695GFP cells subjected to control (top) or GSAP (bottom) siRNA treatments. 3D Volume view of the cells with detected APP-GFP trajectories overlaid and color coded according to their temporal order. For each 10-minute 3D time-lapse recording, we collected 100 Z stacks at an interval of 6 seconds. Each Z stack consists nineteen 100-millisecond optical sections of 512×128 pixels at a Z-axis spacing of 1 μm. Scale bar: 10 μm. (b) Comparison on the distribution of 3D trajectory dwell time (τ_3D_) in single N2a-695GFP cells subjected to control siRNA or GSAP siRNA treatments shows no statistically significant difference (p=0.117).(c) Distributions of trajectory mean velocity 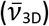 of individual 3D trajectories in N2a-695GFP cells under control or GSAP siRNA treatments shows significant difference between the two groups. (d) Comparison on distributions of total distance traveled (*d*_tot_) of individual 3D trajectories in N2a-695GFP cells under control or GSAP siRNA treatments shows no statistically significant difference (p=0.176). (e) Comparison on distributions of displacement (Δ*d*_3D_) of individual 3D trajectories in N2a-695GFP cells under control or GSAP siRNA treatments shows statistically significant difference. (f) Log-log plot of the mean squared displacement over lag time 〈*r^2^*〉/Δt) as a function of lag time (Δt). In the control siRNA treated cells, diffusion is anomalous around two orders of magnitude in time. Treatment of GSAP siRNA reduces the duration of the anomalous period and an intersection time at around 100 seconds, showing the transition of APP-GFP vesicles from a Trapping to a Trap-Free state. The dashed black line denotes linear fit to the first six data points of the GSAP siRNA group. Whiskers in box plots correspond to one sigma confidence intervals. The distributions were compared using the Mann-Whitney *U* test. ***p < 0.001; ^NS^p > 0.05.

The 3D trajectory dwell times (τ_3D_) of single APP-GFP vesicles were next analyzed to study the dynamic properties in the entire cellular environment (Fig. 7b). Interestingly, unlike the trajectory dwell times acquired by 2D SPT (Fig. 4d), we did not observe significant difference (*p* = 0.117, Mann-Whitney *U* test) of the 3D trajectory dwell times between control group (10 cells from 3 independent experiments, medium τ_3D_ = 36.460 s; IQR 18.158 - 78.405, *n* = 912) and GSAP knockdown group (11 cells from 3 independent experiments, medium τ_3D_ = 36.687 s, IQR 18.266 - 85.606, *n* = 1266). This was expected since trajectory dwell times in 2D SPT are largely limited by the axial detection range of the objective, where the trajectory dwell times generated from mobile pool of APP vesicles are easily underestimated; whereas 3D SPT, although limited in temporal resolution, permits the vesicle tracking covered a much larger range on the vertical scale (up to tenths of microns). Next, we quantified trajectory mean velocity 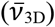 and found that GSAP knockdown results in significant increases in vesicle mobility (medium 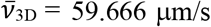, IQR 42.941 - 85.095, *n* = 1266) as compared to control (medium 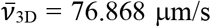, IQR 59.732 – 89.0169, *n* = 912) (Fig. 7c). These results suggested that the effect of GSAP on vascular APP trafficking seems to have more weight on the spatial control than temporal regulation, as no significant differences were found on the trajectory dwell times between the control and GSAP siRNA treated groups.

With this consideration, we next analyzed the spatial properties of vesicle motion by quantitatively measuring the total distance traveled (d_tot_) and the 3D spatial displacement (Δd_3D_) of individual APP-GFP vesicles. Although we found no significant difference in total distance traveled between APP-GFP vesicles subjected to control or GSAP siRNA treatments (*p* = 0.143, Mann-Whitney *U* test; Fig. 7d), vesicles subjected to GSAP siRNA treatment significantly increased 3D spatial displacement as compared to control (\*\*\**p* < 0.001, Mann-Whitney *U* Test; Fig. 7e). The increased velocity of APP vesicles, along with elevated 3D spatial displacement after GSAP knockdown, reflects a preferential decrease of confinement to the surrounding microenvironment. To investigate the underlying sub-diffusion mechanisms further, we fitted our data with the finite trap hierarchy model^53^, which characterizes the obstructed diffusion of individual particles in the biological systems due to barriers, clustering, compartmentation, and/or binding. In the finite trap hierarchy model, the differences between normal diffusion and anomalous sub-diffusion are readily distinguishable by the log-log plot of 〈*γ^2^*〉/Δ*t* versus log Δ*t*, where 〈*γ^2^*〉 is mean square displacement and Δ*t* is lag time. In the log-log plot, anomalous sub-diffusion yields a straight line with a negative slope and the normal diffusion yields a horizontal line. The intersection of both lines defines the crossover time, which represents the transition of single particle diffusion from the trapping state to the trap-free state^53^. Figure 7f shows the log-log plot of APP-GFP vesicles acquired by 4D microcopy. In the control cells, diffusion remains anomalous throughout the entire time, demonstrating that the cellular movement of APP-GFP vesicles stays in the trapping state over the time-course of our measurements. In comparison, APP-GFP vesicles in cells subjected to GSAP knockdown display a crossover to normal diffusion at longer times (Fig. 7f, dashed gray line), indicating that intracellular traps/obstacles attributing to the diffusing APP-GFP vesicles are relieved at longer times after GSAP knockdown.

### GSAP knockdown selectively decreases APP-CTF partitioning in lipid raft

The effect of GSAP on APP trafficking suggested that GSAP may interact with APP. We utilized co-immunoprecipitation assays to determine which GSAP domain interacts with APP. Immunoblot analysis confirmed that GSAP co-precipitated with full-length APP and APP-CTF in HEK293T cells (Fig. 8a, top rows), demonstrating that APP C-terminus is essential for its interaction with GSAP^54^. Given our previous findings that GSAP is processed to release a C-terminal 16 kDa active form (GSAP-16K) ^30^, we asked if GSAP interacts with APP through the GSAP-16K domain. Using the same approach, we found that GSAP-16K co-immunoprecipitated with full-length APP (Fig. 8a, bottom row, left). Furthermore, in accordance with our previously reported results^30^, we also observed that APP-CTF co-immunoprecipitated with GSAP-16K (Fig. 8a, bottom row, right).

Since GSAP interacts with APP, we postulated that GSAP regulates the clustering and partitioning of microenvironments that favor amyloidogenic processing of APP. Amyloidogenic processing of APP is known to mainly occur in the cholesterol-and sphingolipid-enriched membrane microdomains, also known as lipid rafts^55,56^, so we asked whether GSAP resides in lipid rafts, and, if so, whether GSAP knockdown affects APP distribution in lipid rafts. We analyzed the endogenous distribution of GSAP in the lipid raft and non-raft fractions in N2a-695 cells using gradient ultracentrifugation^56^. Lipid raft fractions were identified by the enrichment of raft marker flotillin-2 and fractions located at the bottom of the gradient were labeled as non-raft components. GSAP was specifically detected in the flotillin-2-positive fractions (Fig.8b). In agreement with previous studies, APP-CTF and PS1-NTF were both found to be enriched in the flotillin-2-positive fractions, indicating the accumulation of GSAP, APP-CTF and γ-secretase catalytic component PS1 in lipid rafts where amyloidogenic processing takes place. Considering that GSAP interacts with APP-CTF (Fig. 8a), and that both proteins are enriched in the lipid raft-containing fractions (Fig. 8b), it is strongly suggesting that GSAP regulates lipid raft-dependent amyloidogenic processing of APP-CTF.

**Figure 8.**
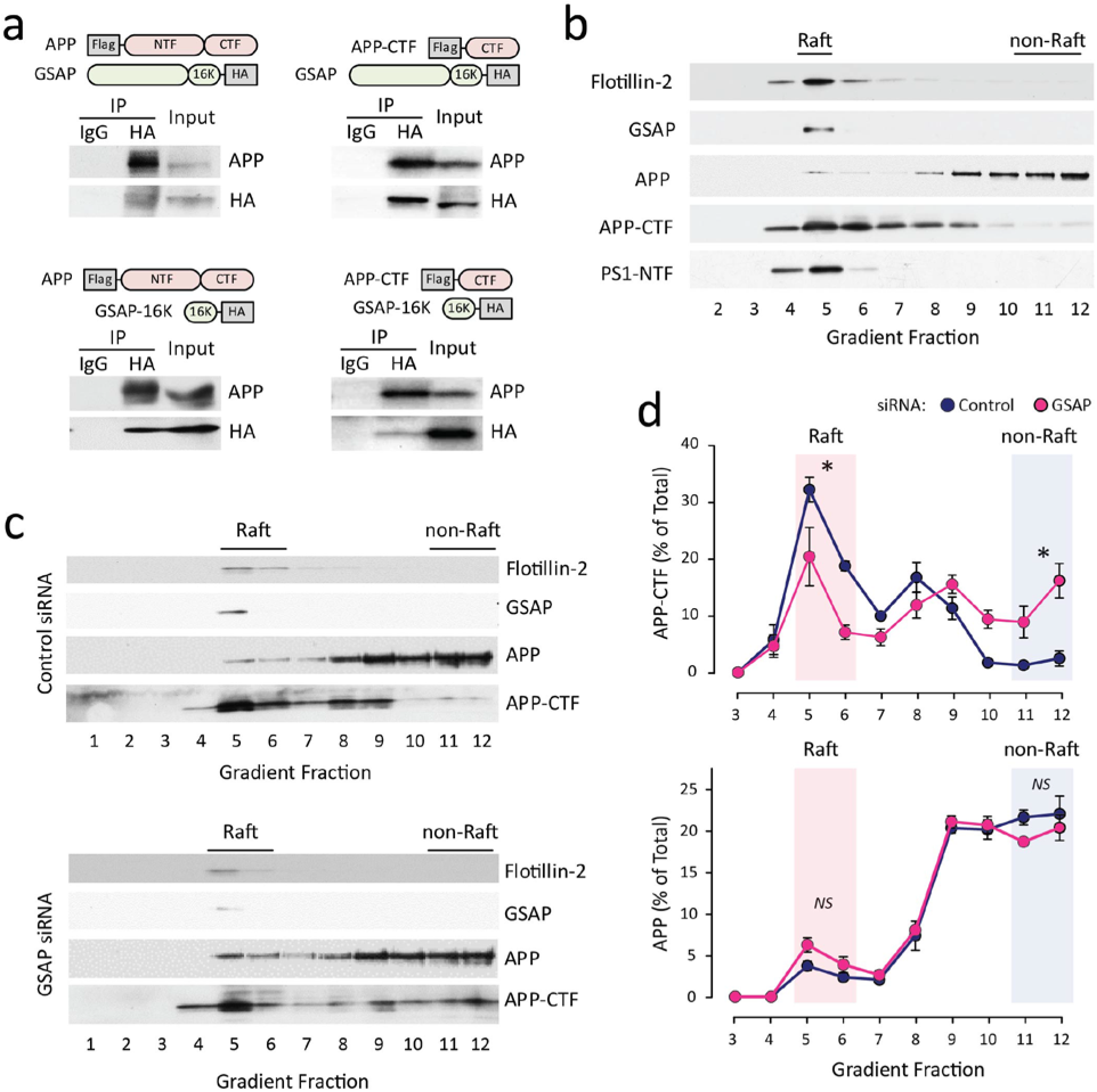
Effect of GSAP knockdown on lipid raft localization of APP and APP-CTF. (a) Representative Immunoblots from co-immunoprecipitation showing the protein-protein interaction between combinations of full-length APP (APP), full length GSAP (GSAP), β-cleaved carboxyl-terminal fragment of APP (APP-CTF), and GSAP c-terminus 16kDa domain (GSAP-16K). Schematic diagrams indicate the plasmids transfected in the HEK293T cells (sketch not to scale). Immunoprecipitates were analyzed by immunoblotting using specific antibodies indicated in the figure. Unspecific IgG were used as the negative control. (b) Representative Immunoblots of lipid raft fractionation showing the enrichment of GSAP in lipid rafts. N2a-695 cells were lysed and subjected to flotation sucrose density gradient centrifugation. Equal volumes of fractions were analyzed by Immunoblotting with indicated antibodies. Flotillin-2 was used as the lipid raft marker. Note that significant amount of GSAP, APP-CTF, and PS1-NTF are present in fractions enriched with Flotillin-2. (c) Representative Immunoblots of lipid raft fractionation showing the effect of GSAP knockdown on lipid raft distribution of APP and APP-CTF. N2a-695 cells were transfected with control or GSAP siRNA then subjected to sucrose density gradient fractionation. Equal volumes of fractions were analyzed by immunoblotting with indicated antibodies. Fraction 5 and 6 were noted as raft fractions and the bottom fractions from 11 to 12 were noted as non-raft fractions. (d) Quantification of the proportion of APP-CTF (top) and APP-full length (bottom) localizing in each fraction. GSAP knockdown decreased the proportion of APP-CTF localizing in lipid rafts and increased the proportion of APP-CTF localizing in non-raft fractions. Data were collected from 3 independent experiments. The distributions were compared using Student’s t-test. *p < 0.05; ^NS^p > 0.05.

Furthermore, we addressed whether GSAP knockdown has any effect on the association of full-length APP and APP-CTF with lipid rafts. GSAP knockdown led to an increase of APP-CTF level in the non-raft fractions located at the bottom of the gradient (Fig. 8c; fraction 10-12). To better quantify the raft localization changes in response to GSAP knockdown, the data from raft and non-raft components (fraction 5-6 and 10-12, respectively) were then combined to reflect the total amount of full-length APP and APP-CTF present within each component (Fig. 8c). The percentage of APP-CTF in lipid rafts was significantly decreased after GSAP knockdown (Δ=23.4 ± 9.2%, P= 0.045). In addition, we also found over a 6-fold increase of APP-CTF in the non-raft fractions (Fig. 8d top). This clearly highlights that the reduction of APP-CTF in lipid raft fractions in response to GSAP knockdown correlated well with an increase of APP-CTF in the non-raft fractions. Such a clear effect of GSAP knockdown on lipid raft-dependent APP-CTF localization raises the question if and how GSAP knockdown also affects the association of APP in lipid raft. Unlike APP-CTF, full-length APP exists mostly in the non-raft fractions and only a small subset of APP is processed in the amyloidogenic pathway ^57,58^. We further evaluated the role of GSAP on lipid raft distribution of full-length APP. Interestingly, the percentage of APP in lipid raft and non-raft fractions were not significantly different regardless of whether N2a-695 cells were treated with control siRNA or GSAP siRNA (Fig. 8d bottom). Together, these findings support the hypothesis that the presence of GSAP favors amyloidogenic processing in lipid rafts, leading to enhanced γ-cleavage^31^.

## Discussion

In this study, we investigated the heterogeneity of vesicular APP trafficking from cell membrane to its intracellular trafficking routes in the whole cells. In addition to its established effects on γ-secretase activity, we addressed whether GSAP regulates APP trafficking and the mechanism by which this might occur. Prior APP trafficking studies were mostly accomplished using high spatial visualization at a single time point, without taking advantage of coupling spatial and temporal resolution with appropriate signal-to-noise ratio. Coupling TIRF microscopy with comprehensive trajectory analysis strategies, our modeling approach effectively simplified the complexity of vesicular APP trafficking near the plasma membrane into “immobile” and “mobile” pools, depending upon how trajectories of single APP vesicles in response to neighboring corrals *in situ* at cell surface. Importantly, we found that GSAP knockdown significantly decreased the number of immobile APP vesicles without effecting the diffusion coefficient of the immobile pool (Fig. 4a). This suggests that GSAP effects vesicular APP trafficking by retaining APP vesicles in membrane microdomains which favor amyloidogenic processing, rather than by changing the composition of membrane microdomains. It has been shown that surface APP is associated with clathrin-coated vesicles^39^. Our result coincides with prior observations that cell surface trafficking of clathrin-coated vesicles, despite their heterogeneity in dwell time and mobility, is subjected to compartment boundaries^59^.

Interestingly, the regulation of GSAP on vesicular APP trafficking appears across cell membrane into cytoplasm. Combining TIRF imaging, high-speed line-scanning imaging^33^ and 4D microscopy^52^, we found that GSAP knockdown significantly increased the mobility of surface and cytoplasmic APP vesicles as compared to control. The two-state kinetic modeling^51^ fit to the cytoplasmic data revealed reduction of BOUND fraction of APP vesicles after GSAP knockdown (Fig. 6d), and, importantly, the reduction ratio is similar to the data obtained at cell surface (Fig. 4a). This suggests GSAP’s involvement through the endocytic pathway that regulates vesicular APP trafficking^23,24^. Confined diffusion of APP vesicles can result from a wide range of underlying physical interactions including APP clustering^38^, hydrophobic interaction with cholesterol-enriched membrane microdomains^60^, phosphorylation and protein-protein interactions^61–63^, among others. Future studies are required to investigate whether other GSAP-associated proteins are required for the functional APP complex assemblies for Aβ generation.

The nanoscale confinement within APP complex assemblies, as well as the extent of the membrane microdomain enrichment, has been shown to effectively modulate how APP complexes are presented and interact with secretases, which impacts APP processing by the secretases. It was previously shown that cholesterol reduction impacts APP cell surface trafficking^55^, reduces APP partitioning to lipid rafts^55,64,65^, disrupts APP-PS1 interaction^55^, and, as a result, decreases Aβ production^55,64,65^. In contrast, increasing membrane cholesterol facilitates APP clustering to raft microdomains^34^, reduces the mobility of APP vesicles, and increases the percentage of stationary APP vesicles^66^, and consequently leads to increased Aβ production^34,66^. As APP can bind to GSAP that is enriched in the lipid rafts (Fig. 8), it is conceivable that GSAP knockdown reduces Aβ production by regulating the preferential enrichment of raft-preferring APP assemblies in amyloidogenic processing^4^. This result also echoes the previous observation by Schreiber *at al*. that cell surface APP clustering, which is demonstrated by the interchange from free to restricted diffusion, drives APP processing from non-amyloidogenic to amyloidogenic pathways^38^.

In summary, using a sophisticated approach that combines various live-cell microscopy and trajectory modeling methods, we provided experimental evidence that GSAP regulates APP trafficking by immobilizing APP vesicles and promoting the APP complex assemblies in favor of amyloidogenic processing. Given that dysregulation of endosomal APP trafficking is highly associated with sporadic AD ^6^, uncovering mechanisms that regulates switching of APP vesicles between immobile and mobile pools may help identifying new strategies for AD treatment.

## Materials and Methods

### Biochemistry

#### Antibodies

Antibodies used in this study were APP C-terminal antibody (1:4,000, RU369, Proc Natl Acad Sci U S A. 1990 Aug;87(15):6003-6. doi: 10.1073/pnas.87.15.6003.), β-Amyloid antibody 6E10 (1:500, 803001, Biolegend), Phospho-APP (Thr668) Antibody (1:1000, 3823S, Cell Signaling Technology), GSAP antibody (1:1000, AF8037, R&D systems), GSAP antibody (1:1000, PA5-21092, ThermoFisher Scientific), Flag M2 antibody (1:1,000, F3165, Sigma), HA antibody (1:1,000, A190-108A, Bethyl), GAPDH antibody (1:500, sc-365062, Santa Cruz), β-tubulin antibody (1:2000, ab6046, Abcam), and BACE1 antibody (1:1,000, PA1-757, ThermoFisher Scientific).

#### Cell culture and transfection

N2a cells stably expressing APP695 (N2a-695) or GFP-tagged APP695 (N2a-695GFP) and N2a cells were maintained in media containing 50% DMEM and 50% Opti-MEM, supplemented with 5% FBS (Invitrogen) with or without 400 μg/ml geneticin. HEK293T cells were maintained in DMEM with 10% FBS. The siRNAs for GSAP were purchased from Dharmacon (On-TARGET plus J-056450-11 and J-056450-12), and Qiagen (FlexiTube GeneSolution GS212167). The control siRNA was purchased from Dharmacon (On-TARGET plus Control siRNA D-001830-02-05). 50 nM final concentration of siRNA was used for transfection. The plasmids used were mouse full length GSAP with HA tag (EX-Mm30424-M07, Genecopoeia), human GSAP-16k with HA tag (a.a. 733 to a.a. 854 subcloned from full length human GSAP plasmids, EX-Z2830-M07, Genecopoeia), human Arcn1 with Myc and Flag tags (RC210778, Origene), Flag-APP-C99 was a kind gift from Wenjie Luo (Weill Cornell Medical College). siRNA and plasmids were transfected with Lipofectamine 2000 (ThermoFisher Scientific) following manufacturer’s instruction.

#### SDS-PAGE, Western blotting and immunoprecipitation

Cell pellets were washed with PBS, then lysed with either 3% SDS or RIPA lysis buffer (20-118, EMD Millipore) supplemented with protease inhibitor cocktail. Equal amounts of protein were subjected to westernblot analysis using precast polyacrylamide gels (10-20% Tris-HCl or 4-12% Bis-Tris, Bio-Rad). For immunoprecipitation (IP) experiments, cell pellets were washed with PBS before lysing in IP lysis buffer (50 mM Tris-HCl, 150 mM NaCl, 1% CHAPSO, pH=7.4, supplemented with protease inhibitor cocktail, and phosphatase inhibitor cocktail), for 10 min on ice. Lysates were then centrifuged at 13,000 g for 10 min at 4°C. Prior to IP, supernatants were collected and diluted in IP lysis buffer 4 times to reach CHAPSO final concentration at 0.25%. 3 μg Flag antibody or mouse IgG control was incubated with lysates overnight at 4°C with tumbling. The next day, 35 μl protein G magnetic beads (ThermoFisher Scientific) were added into samples for 2 h incubation at 4°C. Protein G magnetic beads were collected and washed for 4 times with lysis buffer containing 0.25% CHAPSO. Immunoprecipitated proteins were eluted with SDS sample buffer supplemented with reducing reagent. Samples were heated at 70 °C for 10 min before subjecting to wester nblot analysis.

#### ELISA

Fresh cell culture media without FBS were replaced and incubated 5 h for collection. Aβ level was determined using the ELISA kit following manufacturer’s instructions (ThermoFisher Scientific). The Aβ level is normalized with total protein level and represented as relative fold change over control.

##### DRM fractionation

DRM fractionation was performed following a published protocol ^67^. Briefly, cells were washed twice in PBS, lysed in DRM lysis buffer (Tris-HCl pH 7.4, 150 mM NaCl, and 5 mM EDTA) containing 0.5% Lubrol WX (17A17, Serva) and protease inhibitor cocktail. Cells were homogenized by passing through a 25-gauge needle 5 times. Homogenized cells were adjusted to 45% final concentration of sucrose (4 ml volume) and placed at the bottom of a 12 ml ultracentrifuge tube. 4 ml of 35% sucrose and 4 ml of 5% sucrose were overlaid successively to form a discontinuous gradient. The gradient was centrifuged for 19 h at 39,000 rpm using the Beckman SW41Ti rotor at 4 °C. Twelve 1-ml fractions were collected from the top of the gradient for westernblot analysis.

##### Surface biotinylation

Surface biotinylation was performed similarly as described with modifications^35^. Briefly, cells were rinsed with ice-cold PBS containing 0.1 mM CaCl2, 1 mM MgCl2 and incubated with 0.4 mg/ml Sulfo-NHS-SS-Biotin (ThermoFisher Scientific) for 20 min at 4 °C. Then cells were washed twice with 100 mM Gycine in PBS and twice with ice-cold PBS. Cells were collected and lysed in RIPA lysis buffer for streptavidin magnetic beads (New England Biolabs) pull down. Bound proteins were eluted with SDS sample buffer supplemented with reducing reagent and subjected to western blot analysis.

##### qPCR

RNA was extracted from cells using Direct-zol RNA miniprep kit (Zymo Research) and TRIzol reagent (ThermoFisher Scientific) as described by the manufacturer. 1 μg of RNA was reverse transcribed using the iScript cDNA Synthesis Kit (Bio-Rad). Quantitative PCR was performed using equal amount of cDNA and FastStart Universal SYBR Green Master (ROX) reagent (Roche Applied Science). Primer sequences used for mouse GSAP were described previously^68^: 5’-TCCAGATCACCAGAGAAG-3’ and 5’-ATCCCACTGAGCCCAAAC-3’, mouse GAPDH 5’-AGGTCGGTGTGAACGGATTTG-3’ and 5’-TGTAGACCATGTAGTTGAGGTCA-3’. Raw Ct values were used to calculate GSAP relative expression fold change over control.

### Microscopy

#### Live-cell confocal microscopy

A Zeiss LSM 710 laser-scanning confocal microscope (Carl Zeiss Microscopy, Thornwood, NY) equipped with a 25-mW multiline argon laser was used for live-cell imaging. Images were collected using a Zeiss Plan-Apochromat 100×/1.4 numerical aperture (NA) oil-immersion objective lens. All images were 1024 × 1024 pixels in size and had a 12-bit pixel depth. During imaging, cells were maintained on a heating stage maintained at 37 °C.

#### TIRF microscopy

A custom-made single-molecule TIRF microscope was built on Olympus IX71 inverted microscope for studying live-cell membrane trafficking with high spatial and temporal resolution. The system is equipped with NA 1.50 100X TIRF objectives (UPLAPO100XOHR), a nano-positioning XY linear motorized stage, a 488 nm laser and a millisecond wavelength switching Lambda DG4 excitation light source (Output range: 330 – 700 nm), and a 512 × 512-pixel back-illuminated EMCCD camera (Hamamatsu) with single-photon sensitivity (no binning 70 fps @ full frame; 4X4 binning 1000 fps @ full frame; over 90% max. QE) coupled with a Hamamatsu W-VIEW GEMINI image splitting module for simultaneous dual channel image acquisition. During imaging, cells were maintained on a heating stage maintained at 37 °C. Individual recording of each sample was performed for 60 seconds at a scanning speed of 1 frame/100 milliseconds. Raw data files were extracted to generate stacks of individual 16 bit TIF images for further trajectory analysis.

#### Line-Scanning Confocal Microscopy

Microscopy setup and imaging acquisition were performed as described previously with slight modification^45^. For two-dimensional time-lapse line-scanning confocal imaging of single APP vesicles in living N2a-695GFP cells, images were acquired using a Zeiss 5 Live line-scanning confocal microscope and viewed with a Zeiss 63×/1.4 NA oil-immersion objective lens. The microscope was maintained at 37 °C using a temperature-controlled housing throughout the entire imaging procedures. Line scan images with scan format of 12-bit 512 × 128 pixels were acquired, which gave an *xy* imaging area of 102.4 × 25.6 μm. Individual recording of each sample was performed for 60 seconds at a scanning speed of 1 frame/100 milliseconds. For three-dimensional time-lapse imaging, 0.5 μm z step covering 19 μm was captured every 6 seconds for 10 minutes, and the line-scan format of 12-bit 512 × 128 pixels stayed the same in the entire time course.

#### Particle Trajectory Analyses

For time-lapse TIRF and line-scanning confocal imaging in two-dimensional plane, positions (*x* and *y* coordinates) and trajectories of the single vesicles were determined by Mosaic Particle Tracker plugin^69^. For each vesicle, the resulting fitted particle position data at each time point were exported to a csv file for further analysis in Matlab (The MathWorks) and Spot-On^51^.

The mean-square displacement 〈*r^2^*〉 values for individual trajectories were calculated according to the formula listed below:

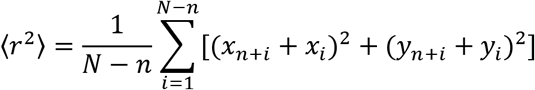

where (*x_n+i_, y_n+i_*) is the position of the particle following a time interval of *nt* (*t* is the time interval between successive measurements) after starting at position (*x_i_*, *y_i_*). *N* is the total number of positions recorded; *n* ranges from 1 to *N*-1. Calculation of mean-square displacement, velocity, diffusion coefficient was processed using custom-written Matlab scripts.

Estimation of the fraction of vesicles in *BOUND* and *FREE* states, and the corresponding diffusion coefficients for each state was determined per individual cell by using the Spot-On software^51^. The jump length histogram was created with a bin width of 0.01 μm and eight time points, four jumps were considered in each track, with a maximum jump length of 3 μm. We fitted a two states model with 0.08 μm^2^/s as maximum *D_BOUND_* and 0.15 μm^2^/s as minimum *D_FREE_* along with default parameters for the “Jump length distribution” and “Model fitting”, except the depth of field was set to 1 μm, corresponding to our line-scanning microscopy.

For 4D microscopy, particle detection and tracking were performed using Imaris 8.2 (Bitplane). Vesicle positions were determined for each frame of the time-lapse z-stack files through the “spot objects” option, and the trajectory paths were constructed using the Brownian motion algorithm. A gap-closing algorithm was also introduced to link trajectory segments resulting from temporary particle disappearance. Trajectory velocity, dwell time, displacement, and mean-squared displacement outputs were used for analysis without any editing.

#### Statistical analyses

All imaging experiments were repeated in at least in three different batches of cultures. Statistical analyses were performed using two-tailed Student’s t-test or Mann–Whitney *U* test using Sigmaplot 13 or PAST statistical analysis package^70^. Nonlinear regression fitting was carried out using Sigmaplot 13.

## Author contributions

Y.-M.L., M.F. and P.G. supervised research; J.C.C. and P.X. designed research; J.C. C., P.X., and E.W. performed research; J.C.C. and P.X. analyzed the data, and J.C.C. wrote the paper.

## Acknowledgments

This work was supported by NIH grant R01AG061350 (to Y.-M.L. and M.F.), the Fisher Center for Alzheimer’s Research Foundation (to P.G. and M.F.), and the JPB Foundation (to Y.-M.L., M.F. and P.G.). Authors also acknowledge the MSK Cancer Center Support Grant/Core Grant (P30 CA008748) and MSK Molecular Cytology Core Facility/Core Grant (P30 CA008748). J.C.C. wish to thank Jesse Aaron and Teng-Leong Chew at HHMI Janelia Research Campus and AIC-HHMI visitation program for the help with Matlab scripts for single-particle tracking of TIRF data. The AIC is jointly supported by the Howard Hughes Medical Institute and the Gordon and Betty Moore Foundation.

