## Supplementary Figures for "GSAP Regulates Amyloid Beta Production through Modulation of Amyloid Precursor Protein Trafficking"

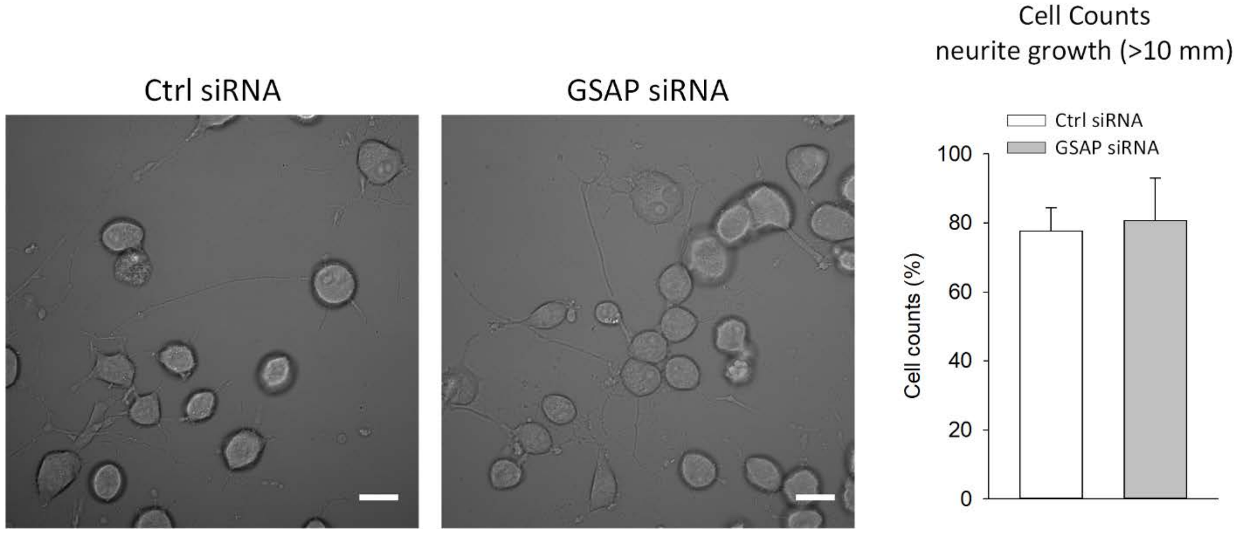


**Suppl. Figure 1**. **GSAP knockdown dose not induce changes in cell morphology and neurite outgrowth.** N2a695 cells were incubated with GSAP siRNA or non-targeting negative control siRNA and allowed to grow for 48 hours prior to observation and morphology analysis of single cells by differential interference contrast (DIC) microscopy. Cells treated with GSAP siRNA or non-targeting control siRNA does not appear markedly different. To quantify the observed morphological changes, we determined the percent of differentiating cells with neurites longer than 10 μm after with either GSAP siRNA or non-targeting control siRNA treatments. Differentiating cells with neurite extensions were prominent and comprised ~80% of the total population in both conditions. Cultures treated with GSAP siRNA vs. non-targeting control siRNA were not significantly different.

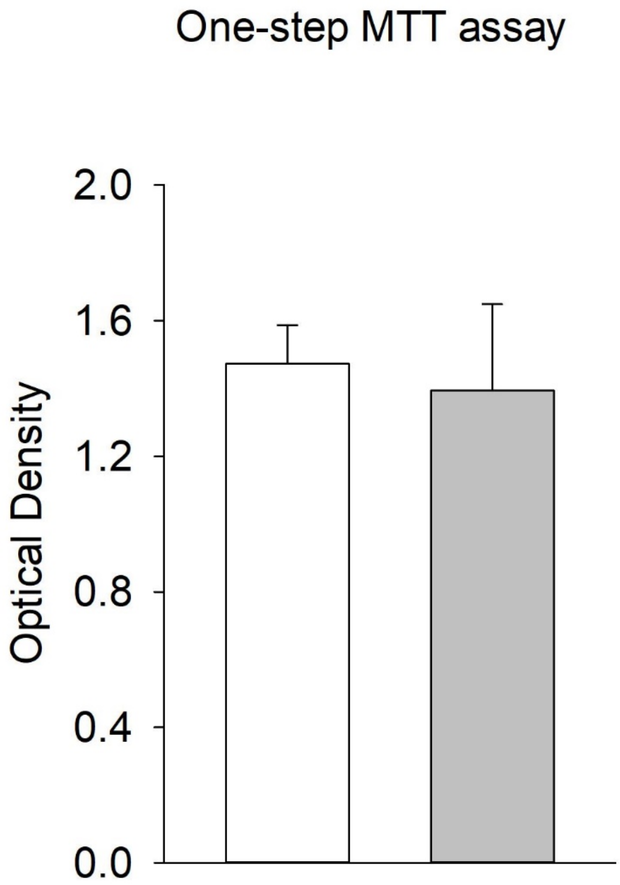


**Suppl. Figure 2. GSAP knockdown dose not induce alteration in cell metabolic activity.** N2a cell cultures were treated with either GSAP siRNA or non-targeting control for 48 hours prior to MTT assay. Measurements were made according to manufacturer's instructions. As shown in the figure, no significant difference in cell metabolic activity was observed between the GSAP siRNA and non-targeting treated conditions.

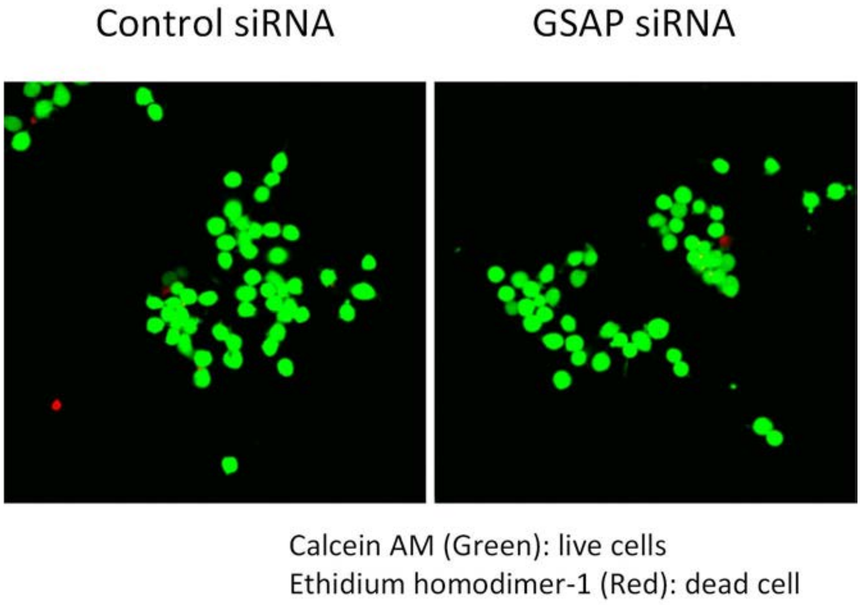


**Suppl. Figure 3. GSAP knockdown has no effect on cell survival.** Assessment of cell viability was performed according to the procedure provided by Sean P. Cregan (ref). N2a cell cultures were treated with either GSAP siRNA or non-targeting control for 48 hours prior to cell viability assay. Cell death/survival is then determined by scoring the fraction of green fluorescent from the cell-permeable substrate Calcein AM (live) and red fluorescent from cell-impermeable dye EthD-1 (dead) neurons using a live-cell confocal microscopy. As shown in the figure, no significant difference in cell death was observed between the GSAP siRNA and non-targeting treated conditions.

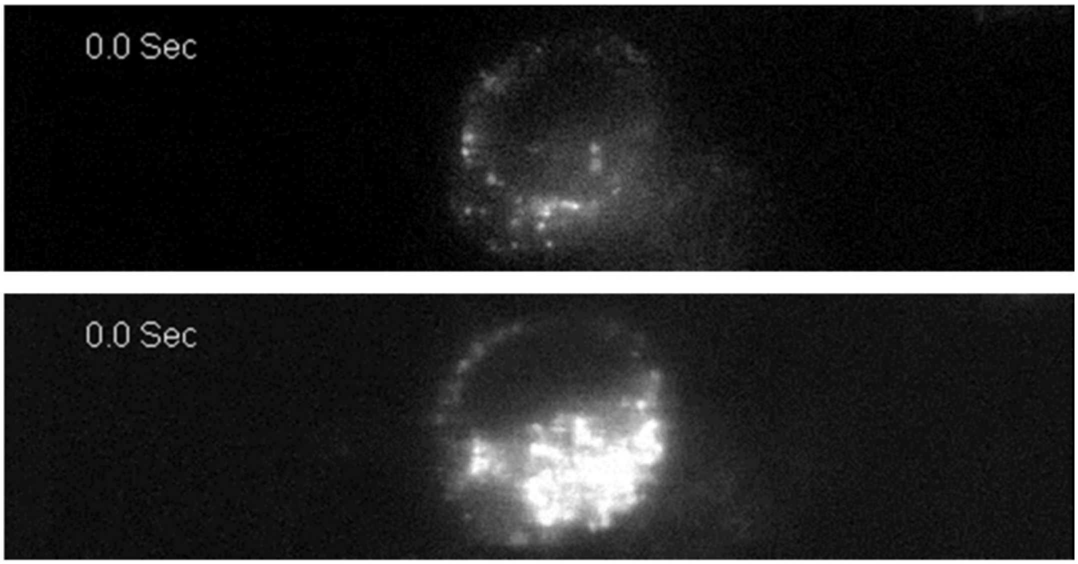


**Suppl. Figure 4. Intracellular APP distribution along the Z-axis.** First frames of the time lapse recording (total 601 frames; frame rate 10 fps) on exact same cell showing the distribution of the APP-GFP fluorescence signal in the cell. A particular vertical level between Golgi and plasma membrane at around 2-3 μm above the coverslip was selected for capturing the dynamic movement of intracellular APP-GFP vesicle (top panel) without high intensity interference from Golgi area (bottom panel).
